# Independent and Interacting Value Systems for Reward and Information in the Human Brain

**DOI:** 10.1101/2020.05.04.075739

**Authors:** I. Cogliati Dezza, A. Cleeremens, W. Alexander

**Affiliations:** Center for Research in Cognition & Neurosciences, ULB Neuroscience Institute, Université Libre de Bruxelles, Brussels, Belgium; Department of Experimental Psychology, Faculty of Brain Sciences, University College London, London, UK; Department of Experimental Psychology, Ghent University, Ghent, Belgium; Center for Complex Systems and Brain Sciences, Florida Atlantic University, USA

## Abstract

Theories of Prefrontal Cortex (PFC) as optimizing *reward* value have been widely deployed to explain its activity in a diverse range of contexts, with substantial empirical support in neuroeconomics and decision neuroscience. Theoretical frameworks of brain function, however, suggest the existence of a second, independent value system for optimizing *information* during decision-making. To date, however, there has been little direct empirical evidence in favor of such frameworks. Here, by using computational modeling, model-based fMRI analysis, and a novel experimental paradigm, we aim at establishing whether independent value systems exist in human PFC. We identify two regions in the human PFC which independently encode distinct value signals. These value signals are then combined in subcortical regions in order to implement choices. Our results provide empirical evidence for PFC as an optimizer of independent value signals during decision-making under realistic scenarios, with potential implications for the interpretation of PFC activity in both healthy and clinical population.

**One Sentence Summary:** Reward and information value are independently optimized in the Human PFC and are then combined in subcortical regions in order to implement choices.

## INTRODUCTION

A general organizational principle of *reward* value computation and comparison in PFC has accrued widespread empirical support in neuroeconomics and decision neuroscience ^1–3^. According to this account, the relative reward value of immediate, easily-obtained, or certain outcomes positively contributes to the net-value of a choice ^4 5^, while delay, difficulty, cost or uncertainty in realizing prospective outcomes negatively contribute to it ^6–8^ (**Figure 1A**). Although substantial empirical evidence supports the interpretation of PFC function as a single distributed system that performs a cost-benefit analysis in order to optimize the net value of rewards ^1–3 9^, other perspectives have suggested the existence of a second, independent value system for optimizing *information* within PFC.

**Figure 1.**
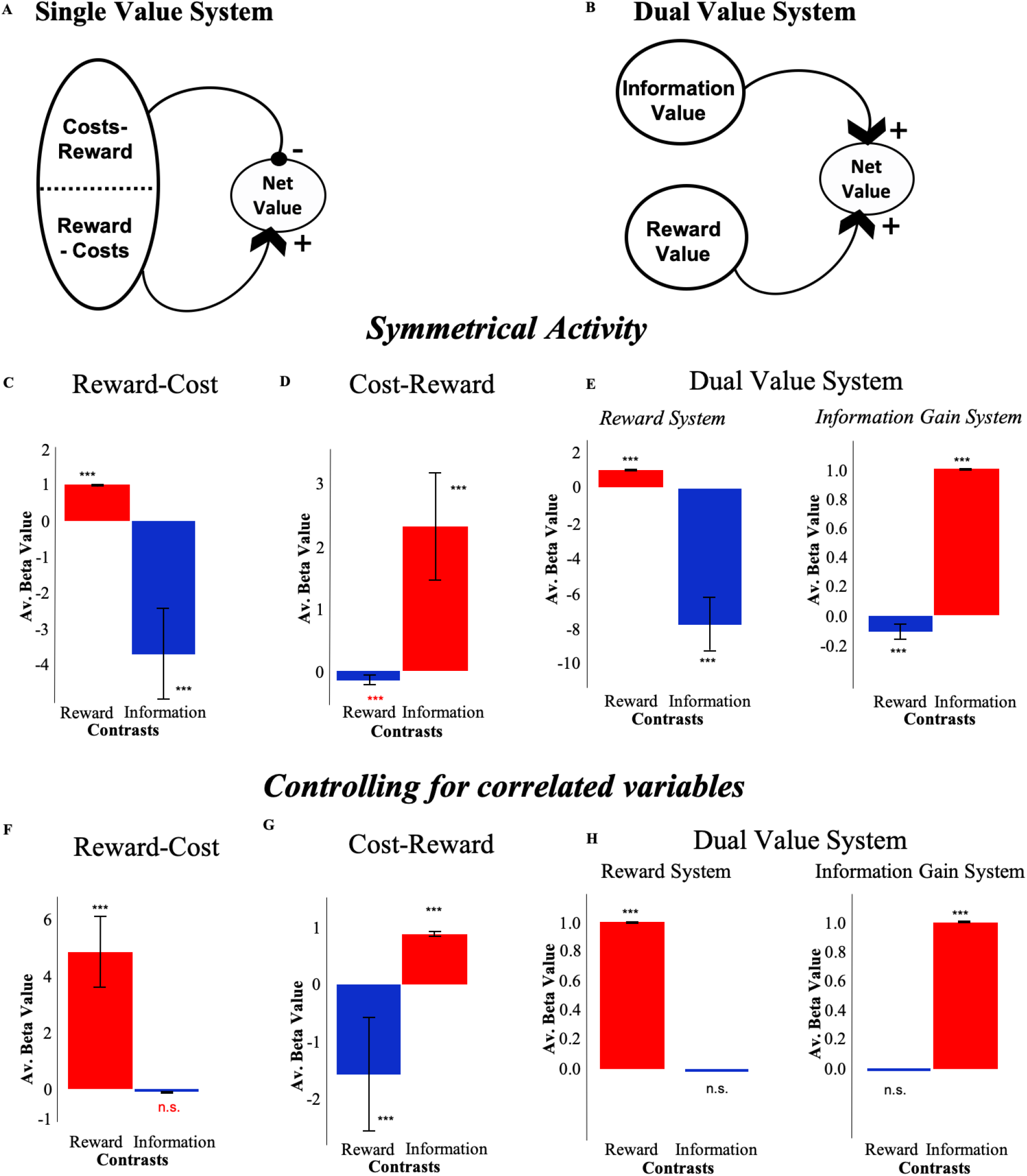
Predictions for single and dual value RL models. **(A)** In the single value system framework costs (negative information value) and rewards interact to produce a net-value estimate, **(B)** while in the dual value system framework information value and reward value are estimated independently. When no controlling for correlations, (**C**) in the single reward value system (Reward-Cost) and (**D**) in the single information value system (Cost-Reward) components representing costs and rewards are represented following an opposing symmetrical pattern of activity. This pattern is also observed in the dual value system, despite the independence of information and reward systems in this framework: optimizing either reward or information gain is associated with decreased activity in the alternate value system, leading to symmetrically opposed activity between the systems **(E)**. When controlling for correlations, symmetrical activity is still observed in the single information value system (**G**), while it disappears in both single reward value system (**F**) and dual value system (**H**).

Unifying theories of brain organization and function propose that information gain plays a similar role as does reward in jointly minimizing surprise ^10–12^, allowing a behaving agent to better anticipate environmental contingencies. Some reinforcement learning (RL) frameworks distinguish extrinsically-motivated (reward-based) behavior from intrinsically-motivated behavior to explain phenomena such as curiosity ^13^, directed exploration ^14^, and play ^15^ in the absence of explicit reward. More broadly, results from machine learning demonstrate that explicitly incorporating information optimization in choice behavior can dramatically improve performance on complex tasks ^16,17^. Altogether these perspectives suggest that information is intrinsically valuable ^18 19^ and positively contributes to the net-value computation ^20 21^ during decision-making (**Figure 1B**). To date, however, direct neural evidence for such a system is lacking. Using computational modeling, model-based fMRI analyses, and a novel experimental paradigm, we aim at establishing whether a dedicated and independent value system for information exists in human PFC. Individuating independent value signals optimized by PFC can offer insight into many disorders characterized by both reward and information-seeking abnormalities such as schizophrenia ^22^, depression ^23^ and addiction ^24^.

Simulations of an RL model which consists of independent value systems ^25^ independently optimizing information and reward demonstrate *how* reward-focused fMRI analyses ^26–30^ may be unsuccessful at identifying an independent information value system as a consequence of correlated activity (**Figure 1**). In such systems, independently optimizing reward and information entails a tradeoff: optimizing reward means *not* optimizing information, and vice-versa. In other words, even if reward and information systems are independent, they are nonetheless (negatively) correlated through behavior ^20^. This correlation is consistent across different decision-making tasks ^20,27,30^ (**Figure 1**; **Supplement**; **Figure S1**), and is also observed in single-value system models (**Figure 1 C,D**). Crucially, the results of our simulations imply that reward-focused univariate fMRI analyses ^26–30^ (which uniquely focus on the reward dimension) may misattribute information value to a system computing costs (diminishing reward value), rather than to an independent information value system.

To test whether a dedicated and independent value system for information exists in human PFC, we focus on two regions, ventromedial PFC (vmPFC) and dorsal Anterior Cingulate Cortex (dACC), which are frequently identified as calculating the positive (vmPFC) and negative (dACC) components of a cost-benefit analysis. Activity observed in vmPFC and dACC frequently exhibits a pattern of symmetric opposition: as dACC activity increases, vmPFC activity decreases-a pattern that holds across a wide range of value-based decision-making contexts, including foraging ^27,30^, risk ^31^, intertemporal ^32,33^ and effort-based choice ^34,35^ (**Table S1**). The variety of contexts in which this pattern is observed suggests a general role for these regions in contributing to the net-value associated with a choice ^1–3^, with vmPFC positively and dACC negatively contributing to the net-value computation. Even for studies in which dACC and vmPFC activity is dissociated ^36 37 38 9 39 40 28^ activity in vmPFC is generally linked to reward value, while activity in dACC is often interpreted as indexing negative or non-rewarding attributes of a choice (including ambiguity ^41^ difficulty ^28^, negative reward value ^35^, cost and effort ^42^). Model simulations further demonstrate that functional opposition between reward and information system in value-based choices may be observed even in absence of a clear symmetric opposition of activity (**Supplement**; **Figure S2**). The interpretation of dACC and vmPFC as opposing one another therefore includes both symmetrically-opposed activity, as well as a more general functional opposition in value-based decision-making.

Here, we used the apparent symmetrical opposition between dACC and vmPFC as a tool to investigate the existence of a dedicated and independent value system for information in human PFC. Simulations of two computational models (**methods)**, one with a single value system (**Figure 1 C,D**) and one with independent value systems (**Figure 1 E**), demonstrate opposed activity for reward value and information gain when these two factors are confounded. Under the single value system model, symmetrically-opposed activity should still be observed after controlling for the confound (**Figure 1 F,G**). In contrast, under the independent value system model, removing the confound between information gain and reward value should abolish symmetric opposition (**Figure 1 H**).

## RESULTS

### Reward and information jointly influence choices

Human participants made sequential choices among 3 decks of cards over 128 games, receiving between 1 and 100 points after each choice (**Figure 2**; **Methods**). The task consisted of two phases (**Figure 2A**): a learning phase (i.e., forced-choice task) in which participants were instructed which deck to select on each trial (**Figure 2B**), and a decision phase (i.e., free-choice task) in which participants made their own choices with the goal of maximizing the total number of points obtained at the end of the experiment (**Figure 2C**). By carefully controlling for reward and information delivered to participants in the learning phase, in this gambling task it is possible to orthogonalize available information and reward delivered to participants in the first free choice trials by clustering choices in specific task conditions ^20^ (**Methods**). To have a better estimate of the neural activity over the overall performance, however, we adopted trial-by-trial fMRI analyses. This introduces information-reward confound in our analysis. Logistic regression of subjects’ behavior on the first trial shows that indeed choices were driven both by the reward (3.22, t(1,19) = 12.4, *p* < 10^−9^) and information (−3.68, t(1,19) = −7.84, *p* < 10^−6^) experienced during the learning phase (**Figure 2D**). In all our analyses, we took care of this confound by using dedicated model-based approaches (**Methods**).

**Figure 2.**
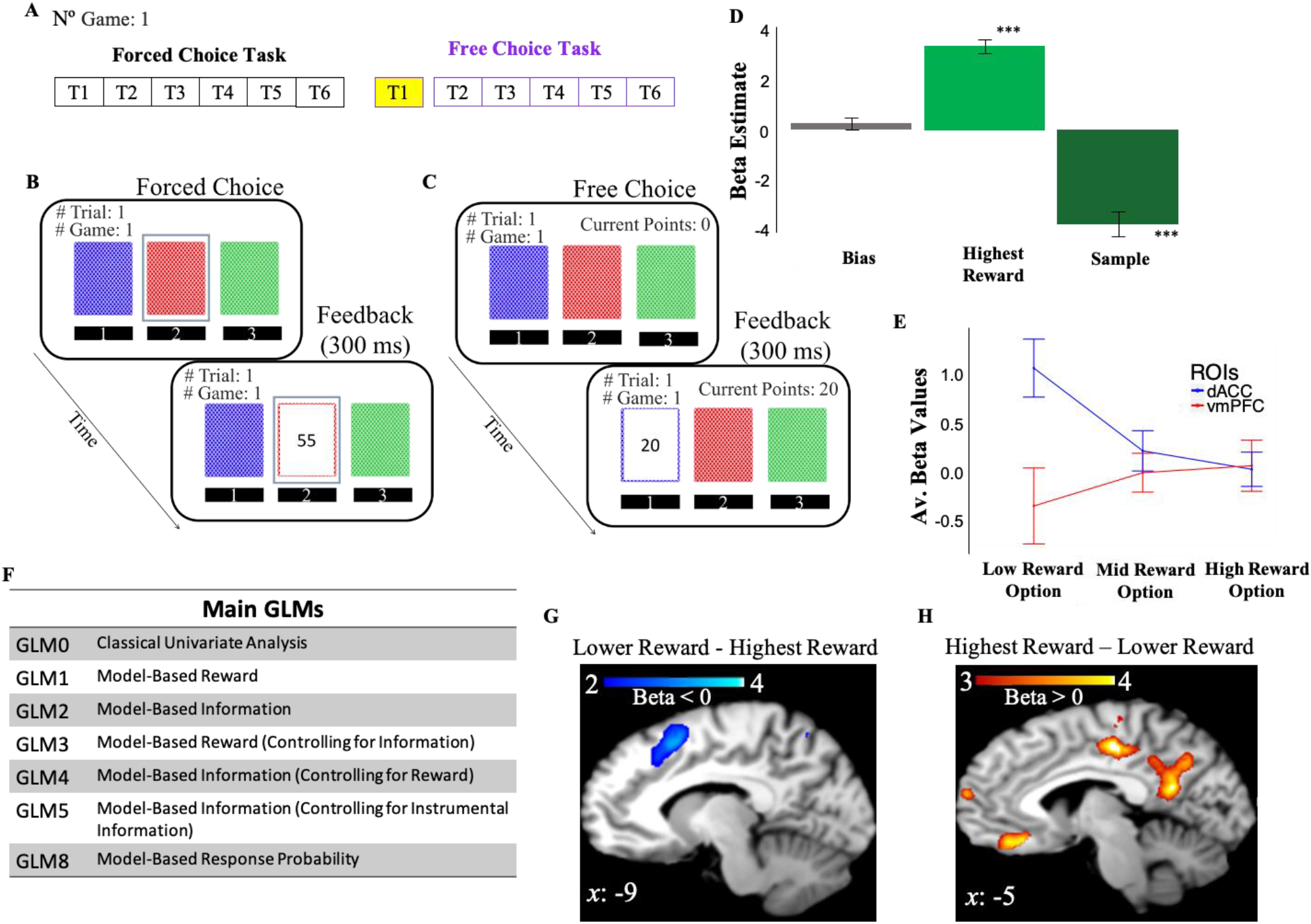
Behavioral Task and Behavior. (**A**) One game of the behavioral task consisted of 6 consecutive forced choice trials and from 1 to 6 free-choice trials. FMRI analyses focused on the first free-choice trial (shown in yellow) in which reward and information were decorrelated. (**B**) In the *forced-choice task* participants chose a pre-selected deck of cards (outlined in blue). (**C**) In the *free-choice task* they were instead free to choose a deck of cards in order to maximize the total number of points. (**D**) Participants’ behavior was driven by both experienced reward and the number of times the options were chosen in previous trials (beta weights from a logistic regression; dependent variable is participants’ exploitative choices). (**E**) DACC and vmPFC activities follow a symmetrical opposite pattern. Activity is split as a function of reward levels (low, mid and high). (**F**) Types of GLMs adopted in the fMRI analysis. (**G**) DACC activity related to selecting the lower reward option. (**H**) VmPFC related to selecting the highest reward option. Activity scale represents z-score.

### The gambling task elicits activity in dACC and vmPFC

We first investigated whether our gambling task elicits dACC and vmPFC activity. We conducted a one sample t-test on the beta weights estimated for *GLM0* which consists of two regressors, one modelling choice onset associated with selection of the highest rewarded options (Highest Reward), and another regressor modelling choice onset associated with lower rewarded options (Lower Reward). This and all subsequent fMRI analyses focus on the time window preceding the first free choices (**Methods**). Results showed that vmPFC activity is positively correlated with the reward associated with the chosen option (*Highest reward − Lower Reward*; FEW p = 0.009, uncorr p = 0.000, voxel extent = 203, peak voxel coordinates (−6, 30, −14), t (19) = 5.48; **Figure 2H**), while dACC/preSMA activity was negatively correlated with the reward of the chosen option (*Lower Reward-Highest Reward*; FEW p = 0.158, uncorr p = 0.014, voxel extent = 87, peak voxel coordinates (−2, 12, 58), t (19) = 4.66; **Figure 2G**). We note that the cluster of activity we identify as “dACC” spans into supplementary motor areas. Many fMRI studies on value-based decision-making reporting similar activity patterns, however, commonly refer to activity around this area as dACC. Additionally, in the Lower Reward – Highest Reward contrast activity did not survive correction for multiple comparisons. This might be due to individual differences in subjective reward value.

We address this issue in the next section by adopting model-based approaches. Here, we repeated the same analysis by selecting a small volume image including the regions that were surviving at uncorr p 0.001. Results show a significant cluster at voxel coordinates (−2, 12, 58) after correcting for multiple comparisons (FEW p = 0.011). Overall, these results indicate that our gambling task elicits activity in dACC and vmPFC and this activity follows a symmetrically opposite pattern (**Figure 2E**).

### Apparent symmetrical activity in dACC and vmPFC as consequence of correlated variables

In the previous section, we showed that our gambling task elicits both dACC and vmPFC activity in a symmetrically opposed pattern. Here, we tested whether 1) this symmetrical activity relates to both reward and information signals and, 2) whether the symmetrically opposite pattern is the product of correlated variables.

We fitted a reinforcement learning (RL) model with information integration ^25^ to participants’ behavior to obtain subjective evaluations of reward and information (**Methods**). Subjective evaluations of reward were computed as relative reward values (*RelReward*; **Methods**) as it has been shown that vmPFC represents the relative expected reward of the chosen option ^28^ (**Supplement**). We additionally considered alternative ways in which vmPFC might compute reward signals. This includes expected reward values, the maximum value of 3 decks, the minimum value of the 3 decks, the reward value variation for the chosen option, the averaged value of the 3 decks, and the value of the chosen option minus the value of the best second option (**Supplement**). These analyses suggested that *RelReward* best describes vmPFC activity. We therefore adopted this computation in the following analyses. Subjective evaluations of information were computed as the prospective information gain for selecting a deck, derived from behavioral fits of our RL model to subject data (*Information Gain*; **Methods**). Alternative computations of information (e.g., the standard deviation of the reward distribution of the chosen option) were highly correlated with this regressor (for most of the subjects r > 0.6) and were therefore not considered any further. RelReward and Information Gain refer to the value associated with the chosen option in the first free-choice trial before the feedback was delivered (**Methods**).

We then regressed RelReward and Information Gain on the BOLD signal recorded on the first free-choice trial of each game. RelReward and Information Gain were used as the only parametric modulators in separate GLMs to identify BOLD activity related to reward (*GLM1*) and to information (*GLM2*) respectively, on the first free-choice trial (**Figure 2F**). Unless otherwise specified, all results for these and subsequent analyses are cluster-corrected with a voxel-wise threshold of 0.001. Activity in vmPFC on the first free-choice trial correlated positively with RelReward (FWE *p* < 0.001, voxel extent = 1698, peak voxel coordinates (8, 28, −6), t (19) = 6.62) (**Figure 3A**) and negatively with Information Gain (FWE *p* < 0.001, voxel extent = 720, peak voxel coordinates (−10, 28, −2), t (19) = 5.36) (**Figure 3B**), while activity in dACC was negatively correlated with RelReward (FWE *p* = 0.001, voxel extent = 321, peak voxel coordinates (6, 24, 40), t (19) = 4.59; **Figure 3A**) and positively with Information Gain (FWE *p* < 0.001, voxel extent = 1441, peak voxel coordinates (8, 30, 50), t (19) = 7.13; **Figure 3B**). A similar symmetrical opposition was observed when we considered expected reward values, instead of relative reward values, for the reward dimension (**Supplement**). Directly contrasting the beta estimates for Information Gain and RelReward in both clusters revealed a symmetrically-opposed pattern of activity in both dimensions (**Figure 3C**). Additionally, we observed that the beta values for GLM1 and GLM2 were negatively correlated across subjects for both the vmPFC cluster (**Figure 3D**) and the dACC cluster (**Figure 3E**).

**Figure 3.**
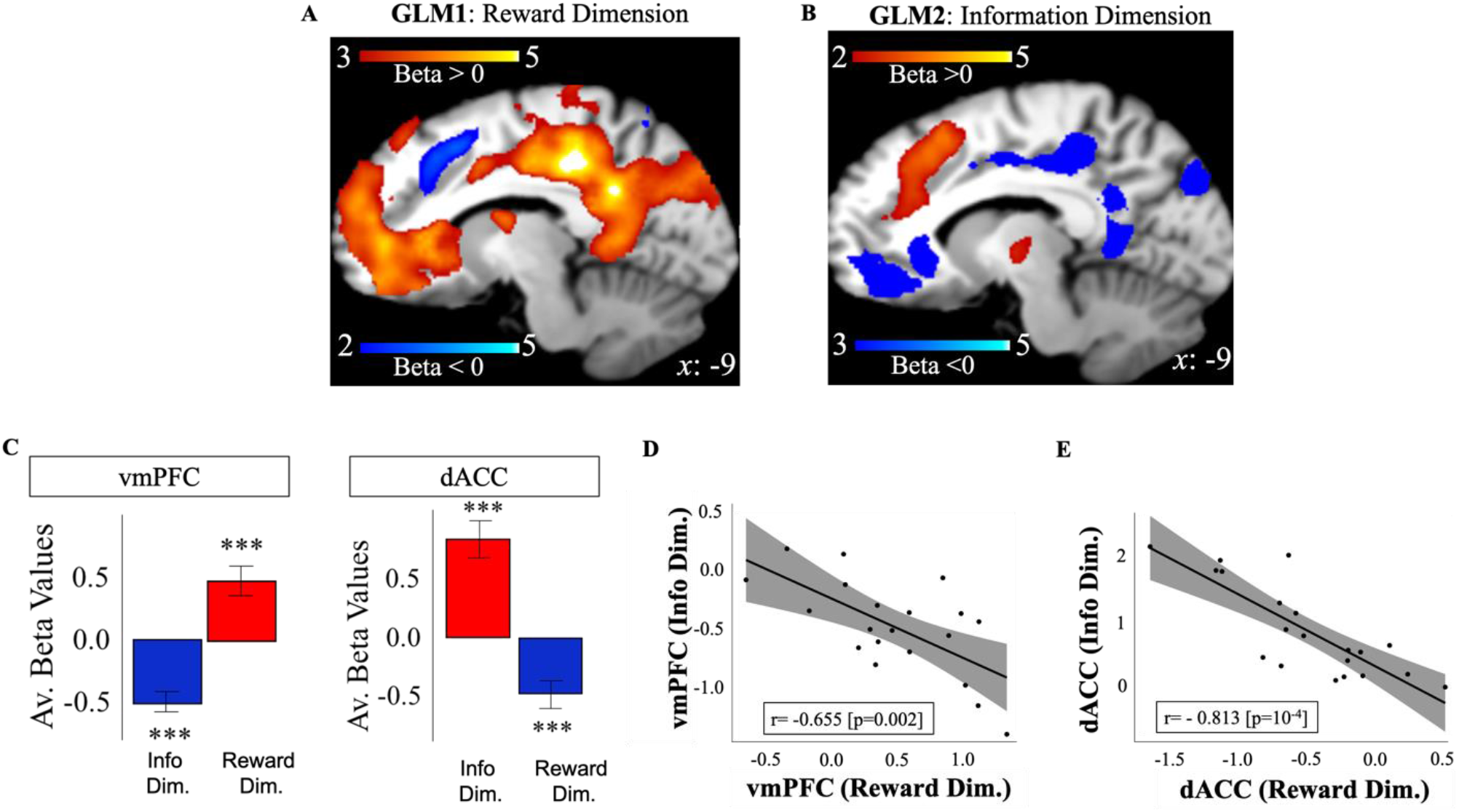
Apperent symmetrical opposition between dACC and vmPFC as consequence of correlated variables. **A**) VMPFC positively correlated with model-based relative reward value for the selected option (in red), while dACC negatively correlated with it (in blue). **B**) DACC (in red) positively correlated with model-based information gain, while vmPFC negatively correlated with it (in blue). Activity scale represents z-score. Averaged BOLD beta estimates **(C)** for each ROI revealed a symmetrically-opposed pattern of activity in both Information and Reward dimensions, in line with model predictions when not controlling for potential correlations (**Figure 1**). BOLD signal estimates for Information Gain and RelReward Value were negatively correlated across subjects for both **(D)** vmPFC and **(E)** dACC ROIs.

The above results showed that the apparent symmetric opposition between dACC and vmPFC is observed along both the information and reward dimension.

### Independent value systems for reward and information

In this section, we repeat our analyses while controlling for possible correlations between information and reward that may underlie our results for GLMs 1 & 2. We investigated the effect of RelReward after controlling for Information Gain (*GLM3*), and the effect of Information Gain after controlling for RelReward (*GLM4*; **Methods**). Activity in vmPFC remained positively correlated with RelReward (FWE *p* < 0.001, voxel extent = 1655, peak voxel coordinates (6, 46, −2), t(19) = 6.56; **Figure 4A**) after controlling for Information Gain in GLM3. In contrast, whereas RelReward was negatively correlated with dACC activity in GLM1, no significant cluster was observed after the removing variance associated with Information Gain in GLM3 as predicted by our model simulations (**Figure 1H**). Similarly, after controlling for the effects of RelReward in GLM4, we observed significant activity in dACC positively correlated with Information Gain (FWE *p* < 0.001, voxel extent = 764, peak voxel coordinates (10, 24, 58), t(19) = 5.89; **Figure 4B**), while we found no correlated activity in vmPFC as was observed in GLM2. A similar pattern of activity was also observed when entering expected reward values, instead of their relative estimates, into GLM4 (**Supplement**).

**Figure 4.**
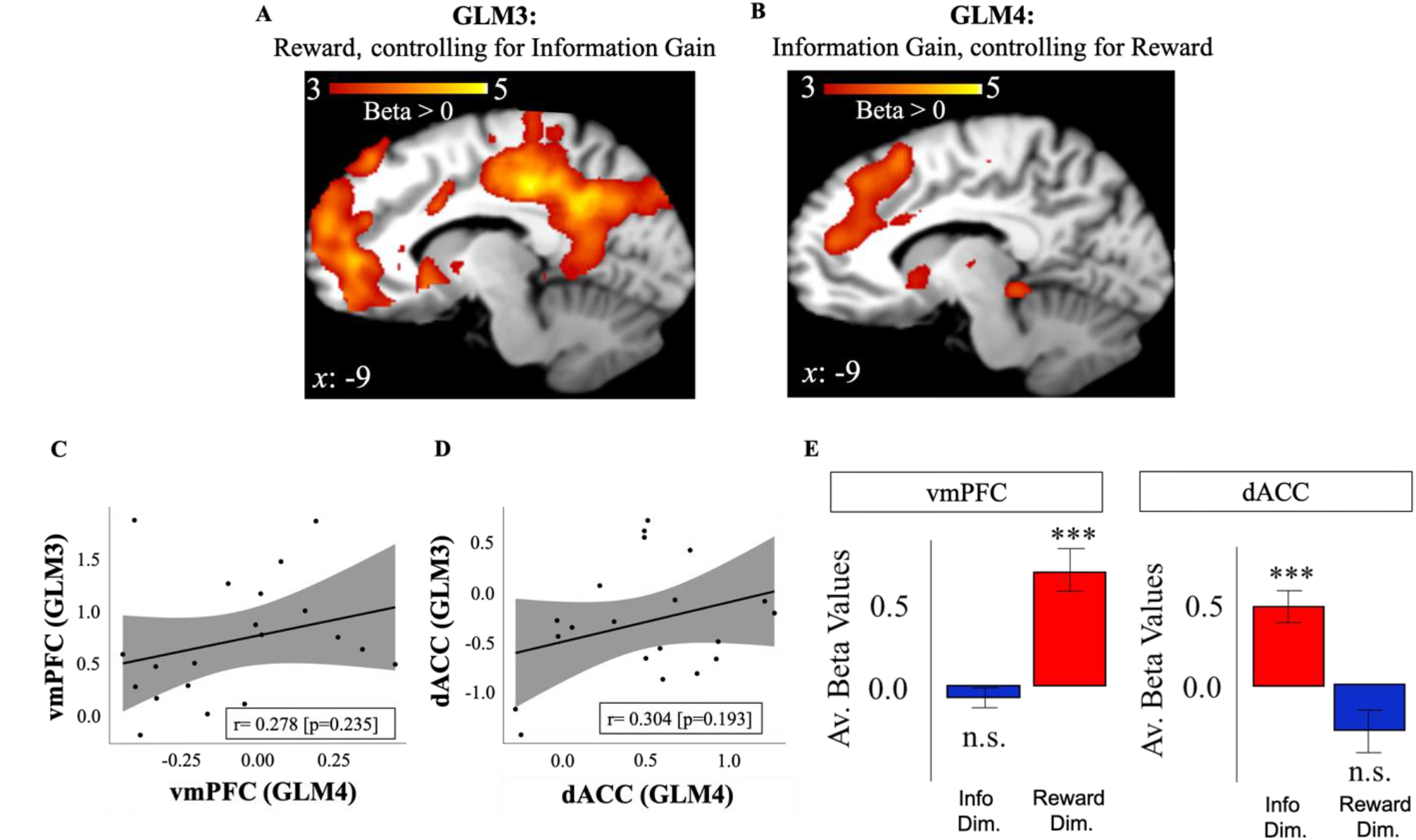
Independent value systems for reward and information in PFC. **A**) After controlling for information effects (GLM3), vmPFC activity (in red) positively correlated with model-based relative reward value (RelReward), while no correlations were observed for dACC. **(B**) After controlling for reward effects (GLM4), dACC activity (in red) positively correlated with model-based information gain (Information Gain), while no correlation was observed for vmPFC. The correlation of BOLD signal estimates between RelReward and Information Gain was no longer observed in either **(C)** vmPFC nor **(D)** dACC, and **(E)** comparison of average BOLD beta values confirms that effects of Information Gain are only observed in dACC, while those of RelReward are observed in vmPFC. “Info Dim” corresponds to the ROIs extracted from GLM4, while “Reward Dim” to the ROIs extracted from GLM3.

Correlations across subjects between the beta estimates for Information Gain (after controlling for RelReward; GLM4) and RelReward (after controlling for Information Gain; GLM3) additionally suggest that activity in vmPFC is specifically related to the relative reward value of the chosen deck (**Figure 4C**) while activity in dACC is specifically related to the information to be gained from the chosen deck (**Figure 4D**) ^43^. Directly contrasting the beta estimates for Information and RelReward in both clusters reveals an asymmetrical pattern of activity in the two dimensions (**Figure 4E)**. These results were replicated after contrasting GLM3 and GLM4 using a paired-t-test (GLM3>GLM4: vmPFC (FWE *p* < 0.001, voxel extent = 467, peak voxel coordinates (−4, 52, 16), t(19) = 5.59); GLM4>GLM3: dACC (FWE *p* < 0.001, voxel extent = 833, peak voxel coordinates (10, 24, 46), t(19) = 5.70); **Supplement**).

To directly test our hypothesis that symmetric opposition between dACC and vmPFC is the product of confounded reward and information signals, we conducted a 3-way ANOVA with ROI (dACC, vmpFC), Value Type (Information Gain, RelReward), Analysis type (confounded{GLM1&2}, no-confounded{GLM3&4}) and we tested the 3-way interaction term. If independent reward and information value signals are encoded in the brain, the two-way interaction (ROI x Value Type) should be significantly modulated by the type of analysis adopted. Results showed a significant 3-way interaction F(1,19) = 19.74, p = 0.0003.

Altogether these findings suggest an independent evaluation of value and the coexistence of two independent value systems for reward and information in human PFC.

### Activity in dACC signals the non-instrumental value of information

In the previous section, we showed that the value of information was independently encoded in dACC. However, in our task different motives may drive participants to search for information ^44^. Information can be sought for its usefulness (i.e., instrumental utility): the acquired information can help with the goal of maximizing points at the end of each game. Alternatively, information can be searched for its non-instrumental value including novelty, curiosity or uncertainty reduction. Here, we tested whether the value of information independently encoded in dACC relates to the instrumental value of information, to its non-instrumental value or to both.

We computed the instrumental value of information (Instrumental Information) by implementing a Bayesian learner and estimating the Bayes optimal long-term value for the chosen option (**Methods**) on the first free-choice trial. We first entered Instrumental Information and Information Gain in a mixed logistic regression predicting first free choices with Instrumental Information and Information Gain as fixed effects, subjects as random intercepts and 0+ Instrumental Information+ Information Gain | subjects as random slope. Choices equal 1 when choosing most informative options, and 0 when choosing options selected 4 times during the forced-choice task. We found a positive effect of Information Gain (beta coefficient = 72.07 ± 16.91 (SE), z = 4.26, *p* = 10^−5^) and a negative effect of Instrumental Information (beta coefficient = −2.25 ± 0.464 (SE), z = −4.85, *p* < 10^−6^) on most informative choices. The percentage of trials in which most informative choices had positive Instrumental Information was ~ 22% suggesting that in most of the trials informative options were not selected based on instrumental utility.

We then entered Instrumental Information and Information Gain as parametric modulators into two independent GLMs. We investigated the effects of Information Gain after controlling for Instrumental Information (*GLM5*) and the effects of Instrumental Information after controlling for Information Gain (*GLM6*; **Methods**). Activity in dACC positively correlated with Information Gain after controlling for Instrumental Information in GLM5 (**Figure 5A**; FWE *p* < 0.001, voxel extent = 1750, peak voxel coordinates (10, 26, 56), t(19) = 8.19). Activity in vmPFC was instead positively correlated with Instrumental Information after controlling for Information Gain in GLM6 (**Figure 5B**; FWE *p* < 0.001, voxel extent = 557, peak voxel coordinates (20, 20, −10), t(19) = 5.62). These results suggest that activity in dACC was strictly related to the non-instrumental value of information, while activity associated with the instrumental value of information was expressed in reward regions consistent with ^45^.

**Figure 5.**
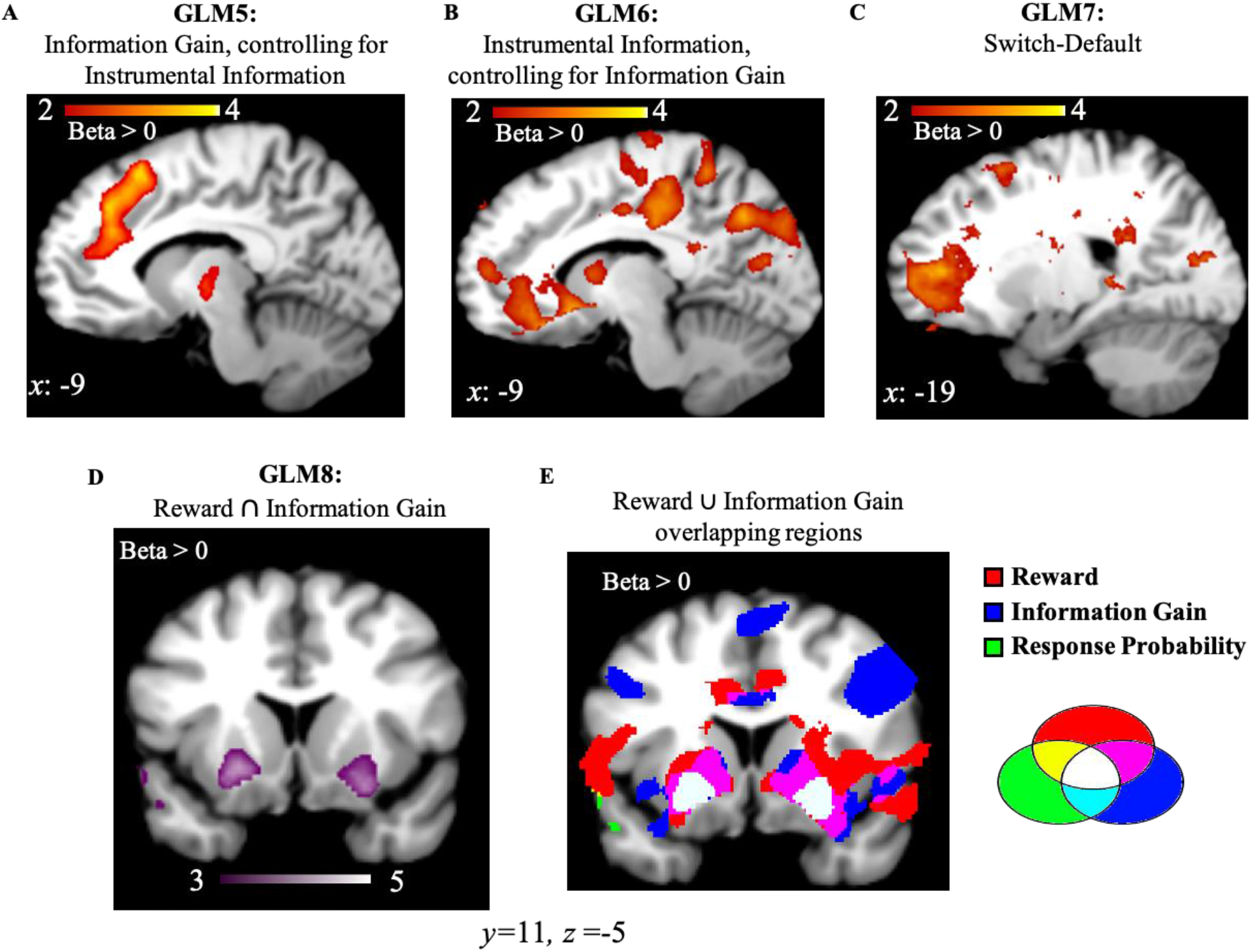
Instrumental Information, Switch strategy, Reward and Information combination in subcortical regions. **A**) Activity in dACC correlated with Information Gain after controlling for the instrumental value of information. **B**) Activity in vmPFC correlated with the instrumental value of information after controlling for Information Gain. **C**) Activity in frontopolar region correlated with Switch (not choosing the most informative options)– Default strategy (choosing most informative options). **G**) Activity in the ventral putamen (striatum region) correlated with response probabilities derived from the RL model and **H**) both RelReward and Information Gain overlap in the striatum region (in white). Activity scale represents z-score.

### Information value and choice difficulty

Activity in dACC has been often associated with task difficulty ^28^ and conflict ^46^. Trials with greater levels of choice difficulty or conflict may lead to prolonged reaction times, and dACC activity may index time on task ^47^ rather than task-related decision variables. In order to rule out the possibility that dACC activity, associated with information value in our task, might instead be driven by time on task, we correlated the standardized estimates of Information Gain with choice reaction times on the first free choice trials. The correlation was run for each subject and correlation coefficients were tested against zeros using a Wilcoxon Signed Test. Overall, correlation coefficients were not significantly different from zero (mean r = 0.031, SE= 0.024; Z= 145; p = 0.1429) suggesting that pursuing an option with higher or lower information gain was not associated with higher or lower choice reaction times as predicted by a choice difficulty or conflict account of dACC function.

### dACC activity does not encode switching strategy

A possible explanation for dACC activity associated with the non-instrumental value of information is that dACC encodes exploration or switching to alternative options away from a default choice. However, in our task the frequency of choosing the most informative option (i.e., exploration) was higher than the frequency of choosing the two other alternatives (in unequal condition, mean = 64.6%, SD = 18 %). Therefore, it is possible that in our task the default strategy was selecting the most informative options, and the switch strategy was selecting less informative (but potentially more rewarding) options. If this is correct, regions associated with exploration or switching strategy (e.g., frontopolar cortex ^36^) should be activated when participants select a non-default option. We conducted a one sample t-test on the beta weights estimated for *GLM7* which consists of a regressor modelling choice onset associated with the most informative options (Default), and another regressor modelling choice onset associated with the other two options (Switch). Activity in frontopolar cortex was positively correlated with Switch (Switch-Default; **Figure 5C**; FWE *p* = 0.019, voxel extent = 148, peak voxel coordinates (−18, 52, 6), t(19) = 5.71). This suggests that adopting a switching strategy in our task was associated with not choosing the most informative options, and not with choosing highly informative options.

### Reward and information signal combine in the striatum

While distinct brain regions independently encode values across different dimensions of the chosen option, these values appear to converge at the level of the basal ganglia. In a final analysis (*GLM8*), we entered choice probabilities derived from the RL model (where reward and information combine into a common option value; eq. 4) as a single parametric modulator, and we observed positively-correlated activity in bilateral ventral putamen (striatum region; right: FWE *p* < 0.01, voxel extent = 238, peak voxel coordinates (22, 16, −6), t(19) = 5.59); left: FWE *p* < 0.01, voxel extent = 583, peak voxel coordinates (−26, 8, −10), t(19) = 5.89; **Figure 4D**). Additionally, ventral putamen overlaps with voxels passing a threshold of p < 0.001 for effects of both RelReward and Information Gain from GLMs 3 & 4 (**Figure 4E**).

## DISCUSSION

Decisions are often influenced by both potential reward and information gains associated with options available in the environment ^25^. Here, we present evidence for dedicated and independent value systems for such decision variables in the human PFC. When correlations between reward and information were taken into account, we found that dACC and vmPFC distinctly encoded information value and relative reward value of the chosen option, respectively. These value signals were then combined in subcortical regions in order to implement choices. These findings are direct empirical evidence for a dedicated information value system in human PFC, independent of reward value. Our finding is in line with a view of human PFC as an optimizer of independent value signals ^10,12,48^.

Our main finding that dACC and vmPFC distinctly encode information gain and relative reward supports theoretical accounts such as active inference ^26 49^ and certain RL models (e.g., upper confidence bound ^47^) which predict independent computations in the brain for information value (epistemic value) and reward value (extrinsic value). Consistent with our findings, the activity of single neurons in the monkey orbitofrontal cortex independently and orthogonally reflects the output of the two value systems ^50^. Therefore, our results may highlight a general coding scheme that the brain adopts during decision-making evaluation.

Our finding that activity in dACC positively correlates with the information value of the chosen option suggests the existence of a dedicated system for information in the human PFC independent of the reward value system. Our control analysis also suggests that dACC encodes the non-instrumental value of information, while the instrumental value of information was expressed in reward regions as previously suggested ^45^. Our results are in line with recent findings in monkey literature that identified a population of neurons in dACC which selectively encodes the non-instrumental value of information ^51^. Additionally, our results are consistent with computational models of PFC which predict that dACC activity can be primarily explained as indexing prospective information about an option independent of reward value ^37,52,53^. DACC has often been associated with conflict ^46^ and uncertainty ^41^, and recent findings suggest that activity in the region corresponds to unsigned prediction errors, or “surprise” ^54^. Our results enhance this perspective by showing that the activity observed in dACC during decision-making can be explained as representing the subjective representation of decision variables (i.e., information value signal) elicited in uncertain or novel environments. It is worth highlighting that other regions might be involved in processing information-related components of the value signal not elicited by our task. In particular, orbitofrontal cortex signals the opportunity to receive knowledge vs. ignorance ^18^ and rostrolateral PFC signals the changes in relative uncertainty associated with the exploration of novel and uncertain environments ^55^. Neural recordings in monkeys also showed an interconnected cortico-basal ganglia network which resolve uncertainty during information-seeking ^51^. Taken together, these findings highlight an intricate and dedicated network for information, independent of reward. Further research is therefore necessary to map the information network in the human brain and understand to what extent this network relies on neural computations so far associated with reward processing (e.g, dopaminergic modulations ^56 57^).

Our finding that vmpFC positively correlates with the relative reward value of the chosen option agrees with previous research that identifies vmPFC as a region involved in value computation and reward processing ^58^. VmPFC appears not only to code reward-related signals ^59 60,61^ but to specifically encode the relative reward value of the chosen option ^62^, in line with the results of our study. We also observed clusters in posterior cingulate cortex which were positively correlated with the relative reward value of the chosen option in a similar fashion as observed for vmPFC, suggesting a role of posterior cingulate in reward processing and exploitative behaviors as previously reported in monkey studies ^63 64^.

Our results further suggest that these independent value systems interact in the striatum, consistent with its hypothesized role in representing expected policies ^65^. The convergence of reward and information signal in the striatum region is also consistent with the identification of basal ganglia as a core mechanism that supports stimulus-response associations in guiding actions ^66^ as well as recent findings demonstrating distinct corticostriatal connectivity for affective and informative properties of a reward signal ^67^. Furthermore, our results are in line with recent evidence on multidimensional value encoding, as opposed to “pure” value encoding, in the striatum ^68 69 70^. Moreover, activity in this region was computed from the softmax probability derived from our RL model, consistent with previous modeling work that identified the basal ganglia as the output of the probability distribution expressed by the softmax function ^71^.

In addition to dACC, we observe activity in additional regions of the cognitive control network which correlate with the information value signal, including bilateral anterior insula cortex and dorsolateral PFC (dlPFC). Activity in these regions is frequently observed in conjunction with dACC activity. Although activity in these additional regions correlates with information value, it is unclear whether they, like dACC, represent information value per se, or instead may represent variables that correlate with information value but were not controlled for in this experiment, e.g., context uncertainty ^53^. Additional work is needed to determine the unique contributions of these regions in signaling information value.

At the same time, our results suggest that the symmetrical opposition between dACC and vmPFC frequently observed in the literature ^27,30 31 32,33 34,35^ can emerge as results of correlated variables. The results of our study are in line with O’Doherty ^72^ who warned of the possibility that, in most neuroeconomics and decision neuroscience studies activity identified as a value signal might instead capture informational signaling of an outcome or particular hidden structure of a decision problem. Recent work has emphasized ecologically-valid tasks for investigating behavior and brain function; while it is critical to characterize the function of brain structures in terms of the behaviors they evolved to support, increased task realism frequently entails a loss of control over experimental variables. Furthermore, we acknowledge that this confound may not emerge in every decision-making studies (e.g., preference-based choice). In those studies, the confound is reflected in subjects’ previous experiences (e.g., the expression of a preference for one type of food over another are consistent, and therefore the subject reliably selects that food type over others on a regular basis). The control of this confound is even more tricky as it is “baked in” by prior experiences rather than learned over the course of an experimental session. Additionally, our results suggest that other decision dimensions involved in most decision-making tasks (e.g., effort and motivation, cost, affective valence, or social interaction) may also be confounded in the same manner. Indeed, symmetrical opposition between dACC and vmPFC has been reported for a wide range of contexts involving decision variables such as effort, delay, and affective valence (**Table S1**). Our findings therefore suggest caution is needed when interpreting findings from such tasks.

Taken together, by showing the existence of independent value systems in the human PFC, this study provides the first empirical evidence in support of a theoretical work aimed at developing a unifying framework for interpreting brain functions. Additionally, this study individuates a dedicated value system for information, independent of reward value. Overall, our results suggest a new perspective on how to look at decision-making processes in the human brain under realistic scenarios, with potential implications for the interpretation of PFC activity in both healthy and clinical populations.

## METHODS

### Participants

Twenty-one right-handed, neurologically healthy young adults were recruited for this study (12 women; aged 19–29 years, mean age = 23.24). Of these, one participant was excluded from the analysis due to problems in the registration of the structural T1 weighted MPRAGE sequence. The sample size was based on previous studies e.g., ^30 33 28^. Participants also presented normal color vision and absence of psychoactive treatment. The entire group belonged to the Belgian Flemish-speaking community. The experiment was approved by the Ethical Committee of the Ghent University Hospital and conducted according to the Declaration of Helsinki. Informed consent was obtained from all participants prior to the experiment.

### Procedure

Participants performed a gambling-task where on each trial choices were made among three decks of cards ^25^ (**Figure 2**). The gambling-task consisted of 128 games. Each game contains two phases: a *forced-choice task* where participants selected options highlighted by the computer for 6 consecutive trials, and a *free-choice task* where participants produced their own choices in order to maximize the total gain obtained at the end of the experiment (from 1 to 6 trials, exponentially inversely distributed such that subjects were most frequently allowed to make 6 free choices). The free choice trial length or horizon was not cued to participants. Therefore, on each game participants were not aware of the free choice horizon. We have already shown that the choice horizon does not affect participants’ choices on this task ^25 73^. In the forced-choice task, participants were forced to either choose each deck 2 times (*equal information condition*), or to choose one deck 4 times, another deck 2 times, and 0 times for the remaining deck (*unequal information condition*). By using this two phase-task, Wilson *et al*. showed that the difference in the number of time each option is sampled and the differences in the mean reward is orthogonalized ^20^ (i.e., options associated with the lowest amount of information were least associated with experienced reward values). In other words, the use of the forced-choice task allows to orthogonalize available information and reward delivered to participants in the first free choice trial. For this reason, the focus of our fMRI analyses is on the first free-choice of each game (resulting in 128 trials for the fMRI analysis). We adopted, however, trial-by-trial fMRI analyses to have a better estimate of neural activity over the overall performance. Therefore, we treated *equal information condition* and *unequal information condition* altogether.

On each trial, the payoff was generated from a Gaussian distribution with a generative mean between 10 and 70 points and standard deviation of 8 points. The generative mean for each deck was set to a base value of either 30 or 50 points and adjusted independently by +/− 0, 4, 12, or 20 points with equal probability, to avoid the possibility that participants might be able to discern the generative mean for a deck after a single observation. The generative mean for each option was stable within a game, but varied across games. Participants’ payoff on each trial ranged between 1 and 100 points and the total number of points was summed and converted into a monetary payoff at the end of the experimental session (0.01 euros every 60 points). Participants were told that during the forced-choice task, they may sample options at different rates, and that the decks of cards did not change during each game, but were replaced by new decks at the beginning of each new game. However, they were not informed of the details of the reward manipulation or of the underlying generative distribution adopted during the experiment. Participants underwent a training session outside the scanner in order to make the task structure familiar to them.

The forced-choice task lasted about 8 sec and was followed by a blank screen, for a variable jittered time window (1 sec–7 sec). The temporal jitter allows to obtain neuroimaging data at the onset of the first-free choice trial and right before the option was selected (decision window). After participants performed the first free-choice trial, a blank screen was again presented for a variable jittered time window (1 sec–6 sec) before the feedback, indicating the number of points earned, was given for 0.5 sec and another blank screen was shown to them for a variable jittered time window. As the first free-choice trial was the main trial of interest for the fMRI analysis, subsequent free-choice trials were not jittered.

### Image acquisition

Data was acquired using a 3T Magnetom Trio MRI scanner (Siemens), with a 32-channel radio-frequency head coil. In an initial scanning sequence, a structural T1 weighted MPRAGE sequence was collected (176 high-resolution slices, TR = 1550 ms, TE = 2.39, slice thickness = 0.9 mm, voxel size = 0.9 x 0.9 x 0.9 mm, FoV = 220 mm, flip angle = 9°). During the behavioral task, functional images were acquired using a T2* weighted EPI sequence (33 slices per volume, TR = 2000 ms, TE = 30 ms, no inter-slice gap, voxel size = 3 x 3 x 3mm, FoV = 192 mm, flip angle = 80°). On average 1500 volumes per participant were collected during the entire task. The task lasted approximately 1h split in 4 runs of about 15 minutes each.

### Behavioral Analysis

#### Expected reward value and information value

To estimate participants’ expected reward value and information value, we adopted a previously implemented version of a reinforcement learning model that learns reward values and information gained about each deck during previous experience - the gamma-knowledge Reinforcement Learning model (gkRL; ^25,73^). This model was already validated for this task and it was better able to explain participants’ behavior compared to other RL models ^4^.

Expected reward values were learned by gkRL adopting on each trial a simple δ learning rule ^74^:

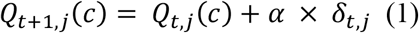

where *Q*_*t,j*_(*c*) is the expected reward value for deck *c* (= Left, Central or Right) at trial *t* and game *j* and *δ*_*t,j*_ = *R*_*t,j*_ (*c*) − *Q*_*t,j*_(*c*) is the *prediction error*, which quantifies the discrepancy between the previous predicted reward values and the actual outcome obtained at trial *t* and game *j*.

Information was computed as follows:

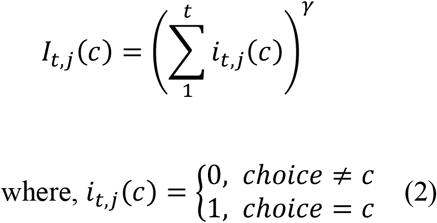

*I*_*t,j*_(*c*), is the amount of information associated with the deck *c* at trial t and game j. *I*_*t,j*_(*c*), is computed by including an exponential term γ that defines the degree of non-linearity in the amount of observations obtained from options after each observation. γ is constrained to be > 0. Each time deck c is selected, *i*_*t,j*_(*c*) takes value of 1, and 0 otherwise. On each trial, the new value of *i*_*t,j*_(*c*) is summed to the previous *i*_*t*−1,1:*j*_(*c*) estimate and the resulting value is elevated to γ, resulting in *I*_*t,j*_(*c*).

Before selecting the appropriate option, gkRL subtracts the information gained *I*_*t,j*_(*c*) from the expected reward value *Q*_*t*+1,*j*_(*c*):

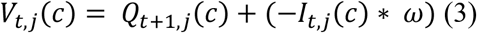

*V*_*t,j*_(*c*) is the final value associated with deck *c*. Here, information accumulated during the past trials scales values *V*_*t,j*_(*c*) so that increasing the number of observations of one option decreases its final value. The scaling is controlled by *ω* which defines the degree new information is integrated in the choice value.

In order to generate choice probabilities based on expected reward and information values (i.e., final choice value), the model uses a softmax choice function ^75^. The softmax rule is expressed as:

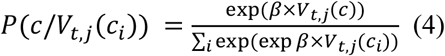

where *β* is the inverse temperature that determines the degree to which choices are directed toward the highest rewarded option. By minimizing the negative log likelihood of *P*(*c*/*V*_*t,j*_(*c*_*i*_)) model parameters α, β, and ω, γ, were estimated for participants’ choices made during the first free-choice trials. The fitting procedure was performed using MATLAB and Statistics Toolbox Release 2020a function *fminsearch* and its accuracy tested using parameter recovery analysis (**Supplement**). Model parameters were then used to compute the value of *Q*_*t*+1,*j*_(*c*) and *I*_*t,j*_(*c*) for each participant. The results of this fit are reported in the table S2.

#### Instrumental value of information

In order to approximate the instrumental utility of options in our task, we turn to Bayesian modeling. In the simplest case, a decision-maker’s choice when confronted with multiple options depends on its beliefs about the relative values of those options. This requires the decision-maker to estimate, based on prior experience, relevant parameters such as the mean value and variance of each option. On one hand, the mean and variance of an option can be estimated through direct experience with that option through repeated sampling. However, subjects may also estimate long-term reward contingencies as well: even if an option has a specific mean reward during one game in our task, subjects may learn an estimate of the range of rewards that options can have even before sampling from any option. Similarly, although subjects may learn an estimate of the variance for a specific option during the forced choice period, over many games subjects may learn that options *in general* have a variance around a specific value.

To model this, we developed a Bayesian learner that estimates, during each game, the probability distribution over reward and variance for each specific option in that game, and, over the entire experiment, estimates the global distribution over mean reward and variance based on observed rewards from all options. A learner’s belief about an option can be modeled as a joint probability distribution over likely values for the mean reward (mu) and standard deviation (sigma). In order to reduce computational demands when conducting forward searches with the model (see below), sigma values were modeled as the integers from 1 to 25 and the range of rewards was modeled in 10 point increments from 5 to 95. Prior to any exposure to the task, the probability distributions over mu and sigma were initialized as a uniform distribution.

To model training received by each subject prior to participation in the experiment, the Bayesian learner was simulated on forced choices from 10 random games generated from the same routine used to generate trials during the experiment. After each choice was displayed, the global probability distribution over mu and sigma was adjusted using Bayesian updating:

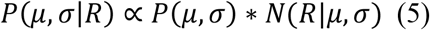

where N() is the probability of observing a reward for a normal distribution with a given mean and variance.

Following the initial training period, the model performed the experiment using games experienced by the subjects themselves, i.e., during the forced choice period, the model made the same choices and observed the same point values seen during the experiment. To model option-specific estimates, the model maintained three probability distributions over µ and σ corresponding to each option, essentially a local instantiation of the global probability distribution described above. The option-specific distributions were reset to the global prior distribution before each new game, and were updated only after an outcome was observed for that option using the same updating rule described in eq. 5.

The Bayesian learner described above learns to estimate the probability distribution over the mean and variance for each option during the forced-choice component of the experiment. If the learner’s only concern in the free-choice phase is to maximize reward for the next choice, it would select the option with the highest expected value. However, in our task, subjects are instructed to maximize their total return for a variable number of trials with the same set of options. In some circumstances, it is better to select from under-sampled decks that may ultimately have a higher value than the current best estimate.

To model this, we implemented a forward tree search algorithm ^76,77^ which considers all choices and possible outcomes (“states”) reachable from the current state, updates the posterior probability distribution for each subsequent state as described above, and repeats this from the new state until a fixed number of steps have been searched. By conducting an exhaustive tree search to a given search depth, it is possible to determine the Bayes optimal choice at the first free-choice trial in our experiment.

In practice, however, it is usually unfeasible to perform an exhaustive search for any but the simplest applications (limited branching factor, limited horizon). In our experiment, the outcome of a choice was an integer from 1 to 100 (# of points), and the model could select from 3 different options, yielding a branching factor of 300. The maximum number of free-choice trials available on a given game was 6, meaning that a full search would consider 3^8 possible states at the terminal leaves of the tree. In order to reduce the time needed to perform a forward tree search of depth 6, we applied a coarse discretization to the possible values of mu and sigma (i.e., sigma values were modeled as the integers from 1 to 25 and the range of rewards was modeled in 10 point increments from 5 to 95). We additionally pruned the search tree during runtime such that any branch that had a probability less than 0.001 of being observed was removed from further consideration.

The value of a state was modeled as the number of points received for reaching that state, plus the maximum expected value of subsequent states that could be reached. Thus, the value of leaf states was simply the expected value of the probability distribution over means (numerically integrated over sigma), while the value of the preceding state was that state’s value plus the maximum expected value of possible leaf states:

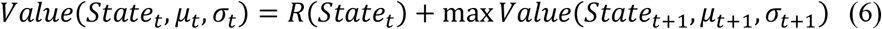

Recursively applying eq. 6 from the leaf states to the first free-choice trial allows us to approximate the Bayes optimal long-term value for each option.

This algorithm was run for each game experienced by a subject during the experiment in order to derive the expected instrumental value (Instrumental Information) for each option following the forced-choice trials specific to that game.

### fMRI analysis

The first 4 volumes of each functional run were discarded to allow for steady-state magnetization. The data were preprocessed with SPM12 (<u>Wellcome Department of Imaging Neuroscience, Institute of Neurology, London, UK</u>). Functional images were motion corrected (by realigning to the first image of the run). The structural T1 image was coregistered to the functional mean image for normalization purposes. Functional images normalized to a standardized (MNI) template (Montreal Neurological Institute) and spatially smoothed with a Gaussian kernel of 8 mm full width half maximum (FWHM).

All the fMRI analyses focus on the time window associated with the onset of the first free-trials prior the choice was actually made (see **Procedure**). The rationale for our model-based analysis of fMRI data is as follows (**Table S3**). First, in order to link participants’ behavior with neural activity, GLM0 was created with a regressor modelling choice onset associated with highest rewarded options (Highest Reward), and another regressor modelling choice onset associated with lower rewarded options (Lower Reward). Activity related to Highest Reward was then subtracted from the activity associated with Lower Reward (giving a value 1 and −1 respectively) at the second level. Next, in order to identify regions with activity related to reward and information, we computed the relative value of the chosen deck (RelReward) and the (negative) value of gkrl model-derived information gained from the chosen option (Information Gain). RelReward was computed by subtracting the average expected reward values for the unchosen decks from the expected reward values of the chosen deck *c* from the gkrl model 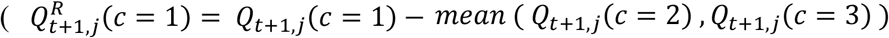. We adopted a standard computation of relative reward values ^28^. It has already been shown that vmPFC represents reward values following the above computation. However, we compare these computations with expected reward values or additional covariates (e.g., chosen-second best option, average of the chosen decks, etc. **Supplement**). Information Gain was computed as −*I*_*t,j*_(*c*). The negative value *I*_*t,j*_(*c*) relates to the information to be gained about each deck by participants. We have already shown that humans represent information value as computed by our model compared to alternative computations when performing the behavioral task adopted in this study ^25^. Next, we entered RelReward and Information Gain as parametric modulators into GLMS with a single regressor modeling the onset of the first free-choice trial as a 0 duration stick function. In GLM1, RelReward was included a single parametric modulator. In GLM2, Information Gain was included as a single parametric modulator. In GLM3, two parametric modulators were included in the order: Information Gain, RelReward. In GLM4, the same two parametric modulators were included, with the order reversed i.e., RelReward, Information Gain. The intent of GLMs 3&4 was to allow us to investigate the effects of the 2^nd^ parametric modulator after accounting for variance that can be explained by the 1^st^ parametric modulator. In SPM12, this is accomplished by enabling modulator orthogonalization (<u>Wellcome Department of Imaging Neuroscience, Institute of Neurology, London, UK</u>). To determine the regions associated with Reward and Information Gain, beta weights for the first (single modulator GLMS) or second (two modulator GLMS) parametric modulators were entered into a 2^nd^ level (random effects) paired-sample t-test. In order to determine activity related to the combination of information and reward value, GLM8 was created with the softmax probability of the chosen option (*P*(*c*/*V*_*t,j*_(*c*_*i*_)) modelling the onsets of first free-choices.

Additional GLMs where then created for the control analyses: GLM1bis with expected reward value (ExpReward) as single parametric modulator; GLM4bis with two parametric modulators ExpReward and Information Gain; GLM5 and 6 with Instrumental Information and Information Gain as parametric modulators; GLM7 which comprises of a regressor modelling choice onset associated with the most informative options (Default), and another regressor modelling choice onset associated with the other two options (Switch); GLM9 and 10 with ExpReward and RelReward as parametric modulators; and GLM11 where the maximum value of 3 decks (Max Value), the minimum value of the 3 decks (Min Value), the reward value variation for the chosen option (Standard Deviation), the averaged value of the 3 decks (Averaged Value), the value of the chosen option minus the value of the best second option (Chosen-Second) and RelReward compete for variance.

Activity for these GLMs are reported either in the main text or in **Table S4**.

In order to denoise the fMRI signal, 24 nuisance motion regressors were added to the GLMs where the standard realignment parameters were non-linearly expanded incorporating their temporal derivatives and the corresponding squared regressors ^78^. Furthermore, in GLMS with two parametric modulators, regressors were standardized to avoid the possibility that parameter estimates were affected by different scaling of the models’ regressors alongside with the variance they might explain ^79^. During the second level analyses, we corrected for multiple comparisons in order to avoid the false positive risk ^80^. We corrected at cluster level using both FDR and FEW. Both corrections gave similar statistical results therefore we reported only FEW corrections.

## Acknowledgments

funded by F.R.S.-fNRS (I.C.D.), FWO-Flanders Odysseus II Award #G.OC44.13N (W.A.) and A.C. was partly supported by an Advanced Grant (RADICAL) from the European Research Council.

## Author Contribution

I.C.D. and W.A. designed and carried out the experiment and discussed the computational modelling and fmri analysis. I.C.D. performed the fmri analysis and the model analysis. I.C.D. and W.A. discussed and interpreted the data. I.C.D, A.C. and W.A. wrote the manuscript.

## Supplementary Material

Supplementary text and Materials and Methods, Figures S1–3, Tables S1-S5, References (*1-6*) accompanies this paper (bottom of this document).

## Supplementary Materials for

Distinct Value Systems for Reward and Information in Human Prefrontal Cortex

## SUPPLEMENTARY TEXT AND RESULTS

### Symmetrical activity in dual and single-value system

#### Dual-value system simulations

To investigate whether correlations between information and reward are expressed in the activity of a dual-value system, we ran 63 simulations of the gkRL model on our gambling-task. The model parameters were selected in the range of those estimated in our sample. We computed Information Gain (− *I*_*t,j*_(*c*)) and RelReward (R*Q*_*t*+1,*j*_(*c*)) in order to simulate the activity associated with the information system and the reward system. Next, we computed model activity in each contrast. To simulate activity of the reward system in the reward contrast we ran a linear regression predicting RelReward with RelReward as independent variable, while in the information contrast we ran a linear regression predicting RelReward with Information Gain as independent variable. To simulate activity of the information system in the reward contrast we ran a linear regression predicting Information Gain with RelReward as independent variable, while in the information contrast we ran a linear regression predicting Information Gain with Information Gain as independent variable. We averaged the beta coefficients across simulations for each contrast and system and we ran a one-sample t test against zeros. As reported in **Figure 1**, averaged beta estimates in the reward and the information contrast were significantly different from zero in both value systems. These results suggest that, even if reward and information are represented by distinct and independent systems, reward and information signals influence activity in a opposite manner.

#### Single-value system simulations

In order to arrive at the overall value of an option, its subjective costs and benefits are weighed against each other. It is generally assumed that the subjective value of an option is modulated by factors that negatively impact the reward value, e.g., the level of effort required, decreases in the probability of obtaining the reward, or longer delays before a reward is received. Similarly, subjective costs may be modulated by the level of reward on offer: discounting rates for effortful, probabilistic or delayed rewards are observed to vary as a function of reward level. In short, subjective costs and benefits are not independent of one another.

We simulated a single-value RL model on our gambling task in which cost was modeled as the uncertainty for selecting an option, consistent with uncertainty aversion observed in human decision-making, and additionally modulated by the relative reward of an option :

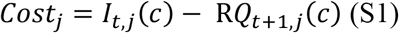

Although uncertainty could be modeled in other ways (e.g., standard deviation of the outcome prediction for an option), for simplicity we adopt the same terms used in the gkrl model. For similar reasons, we model benefits as symmetrical to costs:

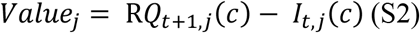

The assumption of a perfectly symmetrical value representation embodied in eq. S2 is likely incorrect – decision-makers are frequently observed to overweight losses relative to gains, for example. Our rationale for modeling a single-value system in this way is to demonstrate as directly as possible how such a system responds to our analysis. Without accounting for the confound between reward and uncertainty/information, the single-value system account makes the same predictions as the dual-system account.

### Accounting for information and reward confound

To account for potential correlations between RelReward and Information Gain we ran the same set of analyses explained above. However, we orthogonalized both RelReward and Information Gain using serial orthogonalization ^1^ to account for potential correlations between these two variables. Next, we simulate activity for the reward system in the information contrast by entering the orthogonalized Information Gain as independent t variable, and to simulate activity for the information system in the reward contrast we entered the orthogonalized RelReward. As shown in **Figure 1**, the symmetrical pattern of activity disappeared.

The same procedure was conducted for the activity recorded from the single-value system model. In contrast to the dual-value system simulations, serial orthogonalization of reward and uncertainty/information *does not* eliminate effects for the alternate decision variable, i.e. we continued to observe the symmetrical pattern of activity for the single-value RL model.

### Information and reward confound in dual-value system generalizes to other decision-making tasks

Here, we tested whether the symmetrical opposition in the dual-value system can be generalized to other sequential decision-making tasks already published in the literature ^2,3^. As in previous versions^2,3^, two cards are displayed on each game and their reward magnitude is visible to the agent. The model has to decide either to engage, which will lead to an economic decision between the two cards (engage cards), or to forage, which will lead to sample alternatives options from the back-up cards. The model has access to the reward magnitude of the back-up cards. Choosing to forage is associated with a cost (ranging between 0 and 3 points). We presented the agent with two conditions: High Information and Low Information. In High Information, half of the back-up cards had lower values than those of the engaged options while the other half had lower values. This condition has maximal uncertainty, i.e., the mean value of new cards obtained through foraging was equally likely to be higher or lower than the mean value of the current cards. Therefore, if the agent decides to forage it has no information on the actual value of the card that will be selected. In the Low Information condition, all back-up cards could have higher or lower values with respect to the engage options. This condition has minimum uncertainty, since the mean value of new cards was guaranteed to be either higher or lower than the current mean card value. The task lasts 135 trials. On each trial, the model computes the value of foraging (i.e., the mean reward of the back-up cards minus the cost of foraging plus the uncertainty associated with the back-up cards: mean (Reward back-up cards) − cost + sd (Reward back-up cards)) and the value of engaging (i.e., the mean reward of engage cards). The model computes decision policies by entering both values into a softmax function. We classified model’s choices as HighReward (when choosing the option-forage or engage-associated with the highest mean reward), LowReward (otherwise); HighInfoGain (when choosing forage in High Information condition) and LowInfoGain (otherwise). Subsequently, we computed Information Gain as the value of foraging and Reward as the value of engaging in order to simulate neural activity associated with the reward system and information system. We then ran a first level analysis over the averaged value of engaging (value of foraging) in the HighReward choices minus the average of reward values in the LowReward choices (Reward Contrast); and the averaged value of engaging (value of foraging) in the HighInfoGain trials minus the averaged value of engage (value of foraging) in LowInfoGain trials (Information Contrast). As already shown with the simulation of our gambling task, activity associated with Reward and Information Contrast are correlated in both value systems and both the reward dimension and information dimension are represented in a symmetrically opposite manner within the two systems (**Figure S1**). These results demonstrate that the confound in the representation of reward and information with value systems can be generalized to other sequential decision-making tasks.

### Information and reward system opposition in value-based choice in absence of symmetrical opposition

Here, we show how functional opposition between reward and information systems in value-based choices may be observed even in absence of clear symmetric opposition of activity, we simulated an effort-based environment where rewards could be obtained only after exerting effort. In many effort-based paradigms^4^, subjects must choose between a small, default reward that requires little effort to obtain, or a larger reward that requires greater effort, and consequently a chance of failing to perform the task adequately and not receiving a reward. We adapted our RL model in order to simulate choices made by an agent performing this task. In this implementation, the information value is equal to the entropy (−p*log(p); where p is the probability of successfully performing the task) resulting in the following value function:

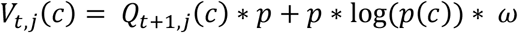

We simulated the model across different ranges of effort and rewards. While the probability of the model selecting the non-default option decreased with effort level (**Figure S2A**) and increased with relative reward value (**Figure S2B**), consistent with research in this area, there was no correlation between the relative reward value and effort level (**Figure S2C**). Finally, for the range of effort levels included in this simulation, the level of effort correlated with the information value signal (**Figure S2D**). This result suggests that even when activity indicating effort and relative reward can be dissociated in value-based decision-tasks, the interpretation that the regions serve functionally opposed roles may be misguided.

### Logistic regression of subjects’ behavior

In order to investigate participants’ behavior during the scanning session, we performed a logistic regression for each participant over exploitative choices against the following normalized variables: highest experienced reward (Highest Reward) and number of samples for the highest rewarded option (N° samples). The dependent variable had binary output {exploitative choices =1; non-exploitative choices – or exploration = 0 otherwise}. Exploitative choice trials were classified as those trials in which participants chose the option in the first free-choice trial associated with the highest average of points collected during the forced-choice task of the same game. Beta coefficients were collected for the entire group and a one sample t-test was conducted as shown in the main text to test whether coefficients significantly differed from 0 (**Figure 2D**).

### Parameter recovery, model comparison and model simulations

We performed a parameter recovery analysis to estimate the degree of accuracy of the fitting procedure. To do so, we simulated data from gkRL using the parameters obtained from the fitting procedure (*true parameters*), and we fit the model to those simulated data to obtain the estimated parameters (*fit parameters*). We then ran a correlation for each pair of parameters ^5^. This revealed high correlation coefficients for α (r= 0.8, *p* < 10^−3^), β (r= 0.8, *p* < 10^−3^), ω (r= 0.6, *p* = 0.006) and γ (r= 0.6, *p* = 0.002).

We additionally ran model comparisons to estimate the degree by which gkRL explained choice behavior in our sample compared to a standard RL model-where only reward predictions influence choices. Negative log likelihoods obtained during the fitting procedure were used to compute model evidence (the probability of obtaining the observed data given a particular model). We adopted an approximation to the (log) model evidence, namely the Bayesian Information Criterion (BIC) ^6^ and we compared its estimate across different models (fixed-effect comparison). Additionally, we used a random-effect procedure to perform Bayesian model selection at group level ^7^ where we computed an approximation of model evidence as –BIC/2. Model comparison showed that the gkRL model was better able to explain participants’ choice behavior compared to a standard RL (xpgkRL= 1, BICgkRL= 14354; xpstandardRL =0, BIC standardRL=16990).

We simulated gkRL using the estimated free parameters. We then performed a logistic regression for each simulation predicting gkRL choices (exploitative choices =1; non-exploitative choices – or exploration = 0) with reward and number of samples as fixed effects. Exploitative choice trials were classified as those trials in which the model chose the option in the first free-choice trial associated with the highest average of points collected during the forced-choice task. Logistic regression was fitted for each individual simulation and beta coefficients tested against zero. Reward and number of samples were significantly impacting gkRL choices (both p < 10^−5^) as observed in our human sample.

### Symmetrical opposition is also observed for expected reward values

In GLM1, the reward regressor was defined as the relative reward value of the chosen option (RelReward). We defined reward in this way based on previous research that showed vmPFC activity associated with the relative reward value of the chosen option ^8^. Here, we tested whether the symmetrical pattern was also observed when computing reward as the expected value of the chosen option (*Q*_*t*+1,*j*_(*c*); ExpReward). We computed GLM1bis with ExpReward as a parametric modulator instead of the relative reward value. ExpReward positively correlated with vmPFC (FWE *p* < 0.001, voxel extent = 530, peak voxel coordinates (−10, 34, 14), t (19) = 5.47) and negatively correlated with dACC (FWE *p* = 0.001, voxel extent = 205, peak voxel coordinates (8, 22, 40), t (19) = 5.47) suggesting that the symmetrical pattern was also observed when considering expected rewards and not just their relative estimates.

### Relative reward value computation best describe vmPFC activity compared to expected reward value

Here, we tested whether the reward signal computed in vmPFC was best described by the relative reward value of the chosen option (our RelReward regressor) or by the expected reward value of the chosen option (ExpReward). We computed GLM9 with ExpReward (controlling for RelReward) and GLM10 with RelReward (controlling for ExpReward) as parametric modulators. Activity in vmPFC positively correlated with relative reward after controlling for ExpReward (FWE *p* < 0.001, voxel extent = 1829, peak voxel coordinates (−6, 52, 14), t (19) = 7.21). However, vmPFC activity associated with ExpReward disappeared after controlling for RelReward suggesting that vmPFC encodes the relative reward value of the chosen option as previously suggested ^8^.

### Additional reward covariates

Here, we investigated the role that additional reward covariates can play in modulating vmPFC activity. We individuated 5 additional covariates: the maximum value of 3 decks (Max Value), the minimum value of the 3 decks (Min Value), the reward value variation for the chosen option (Standard Deviation), the averaged value of the 3 decks (Averaged Value), the value of the chosen option minus the value of the best second option (Chosen-Second). We first computed the correlations with Reward (**Table S5**). Standard Deviation and Chosen-Second showed really high correlation with RelReward therefore were not considered any further. For the remaining variables, we entered them as parametric modulators in a GLM11 together with Reward allowing to compete for variance. We then extracted vmPFC ROI from GLM2 (to avoid that reward activity might interfere with ROI identification) and we computed the averaged beta estimates for each covariate. Next, we compared these betas to the betas estimated in vmPFC ROI for RelReward. Results showed that the betas for RelReward were significantly higher than the betas for Max Value (p < 0.05), Min Value (p < 0.05), Averaged Value (p < 10^−3^) suggesting that the majority of the variance in vmPFC was accounted by our RelReward.

### dACC encodes the value of information after controlling for the expected reward value

The analyses reported in the main text clearly suggest that dACC encodes the value of information. However, in GLM4 the activity associated with the value of information was controlled by the variance explained by the relative reward value. It is possible that the variance could only account for the “relativeness” instead of the reward itself. Therefore, we ran an additional analysis to investigate activity associated with Information Gain after controlling for ExpReward (GLM4bis). Results showed that Information Gain was positively correlated with dACC after controlling for ExpReward (FWE *p* < 0.001, voxel extent = 640, peak voxel coordinates (10, 30, 58), t (19) = 6.87) suggesting that whether we control for expected reward or their relative estimates activity associated with Information Gain was still expressed in dACC.

### Dissociable regions for Reward and Information

While our results from GLMs 3 & 4 demonstrate that activity in vmPFC & posterior cingulate is explained by RelReward after controlling for Information Gain, and activity in dACC & anterior insula is explained by Information Gain after controlling for RelReward, these analyses do not allow us to conclude that one set of regions is specific to reward while the other is specific to information (i.e., while we can say the, for example, RelReward is different than 0, and Information Gain is not different than 0, we cannot say RelReward is different than Information Gain). In order to do so, we directly compare the beta weights estimated for RelReward (after orthogonalizing with respect to Information Gain) from GLM3 and the beta weights estimated for Information Gain (orthogonalized with respect to Reward) from GLM4 using a paired-sample t-test. We find clusters of activity in vmPFC (as reported in the main text), posterior cingulate (FWE *p* < 0.001, voxel extent = 1493, peak voxel coordinates (−14, −48, 36), t(19) = 6.02) and putamen (FWE *p* < 0.001, voxel extent = 920, peak voxel coordinates (24, 10, −8), t(19) = 6.13) in which Reward > Information Gain (Figure S3A), indicating that these regions are specifically involved in reward processing, while a significant cluster is observed in dACC (as reported in the main text), right insula (FWE *p* < 0.05, voxel extent = 157, peak voxel coordinates (34, 24, −6), t(19) = 4.89)) and dlPFC (FWE *p* < 0.05, voxel extent = 158, peak voxel coordinates (48, 32, 32), t(19) = 4.89) for (negative) Information Gain > RelReward (Figure S3B), indicating that dACC is specifically involved in representing uncertainty.

---

**Figure S1.**
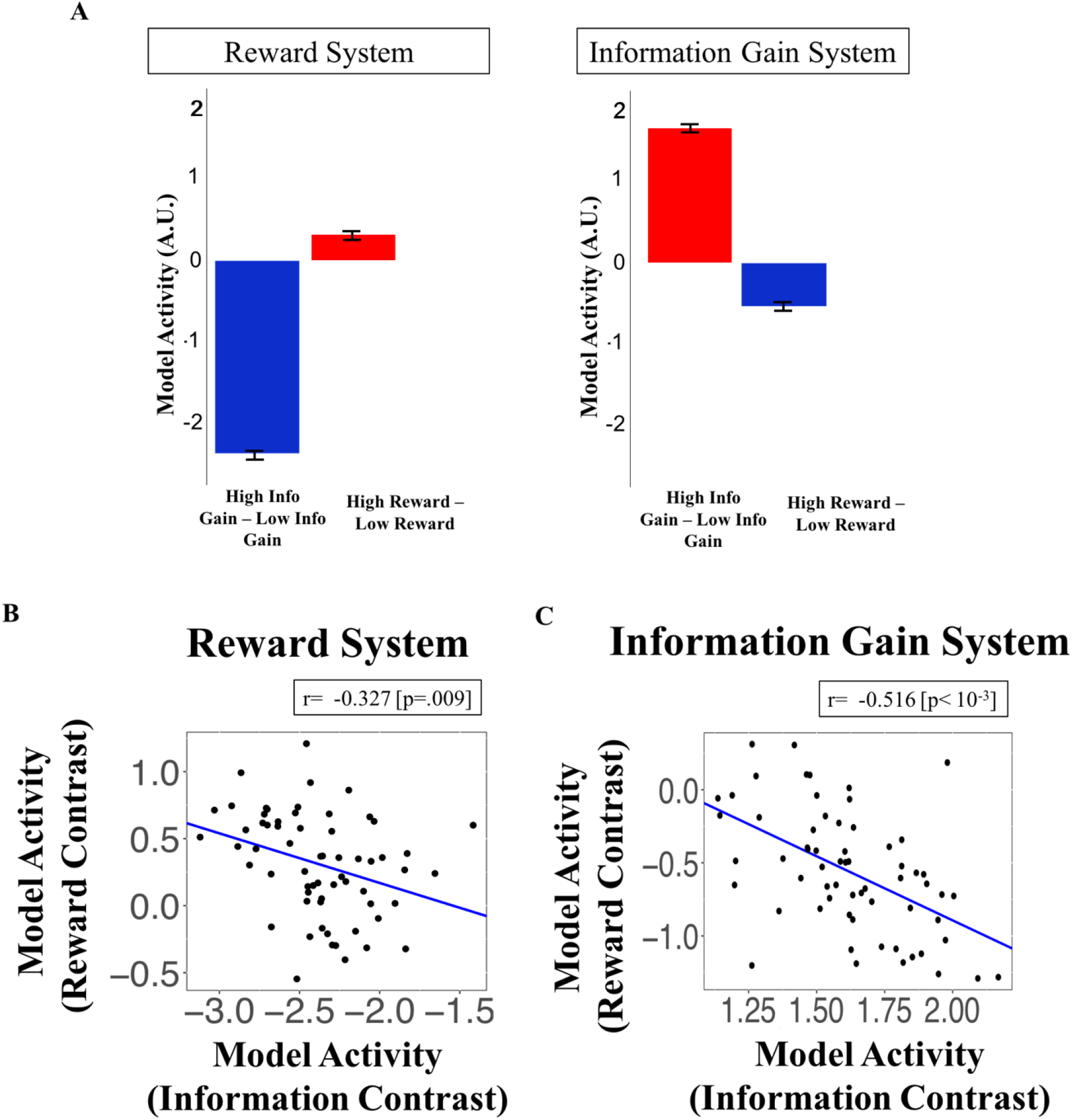
Correlated activity in foraging task. **A)** Simulating a dual value system on the sequential decision-making task adopted by ^3 2^. Despite the independence of information and reward systems, the systems’ activity are correlated: optimizing information is associated with decreased activity in the reward value system, and optimizing reward is associated with decreased activity in the information value system. Activity within the **(B)** reward system and **(C)** the information system is negatively correlated across independent model simulations.

**Figure S2.**
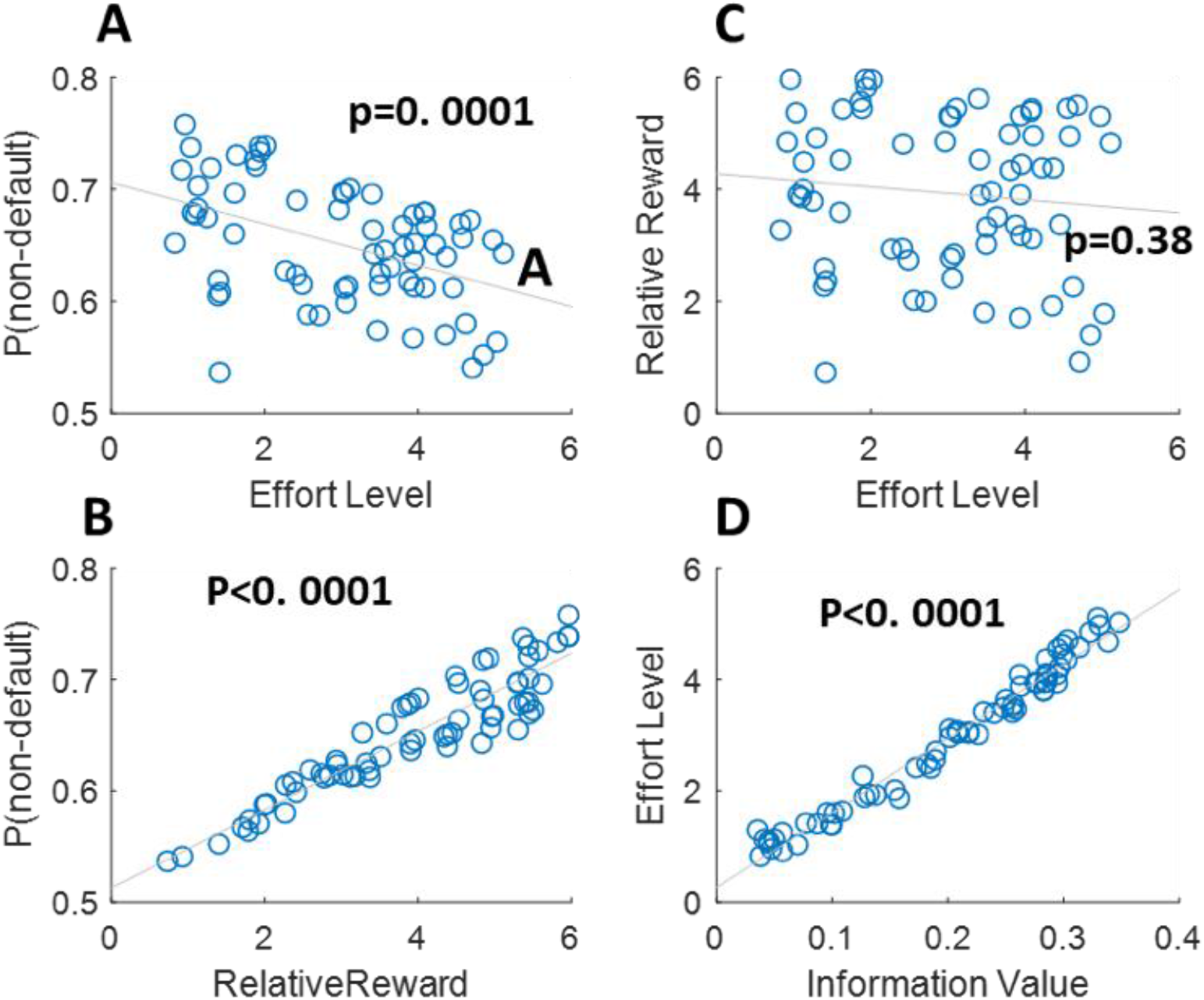
Functional opposition between vmPFC and dACC in absence of symmetrical opposition. Probability of the model selecting the non-default option across effort levels (**A**) and its relative reward values (**B**). Correlation between relative reward value and effort levels (**C**). Correlation between information value and effort levels (**D**).

**Figure S3.**
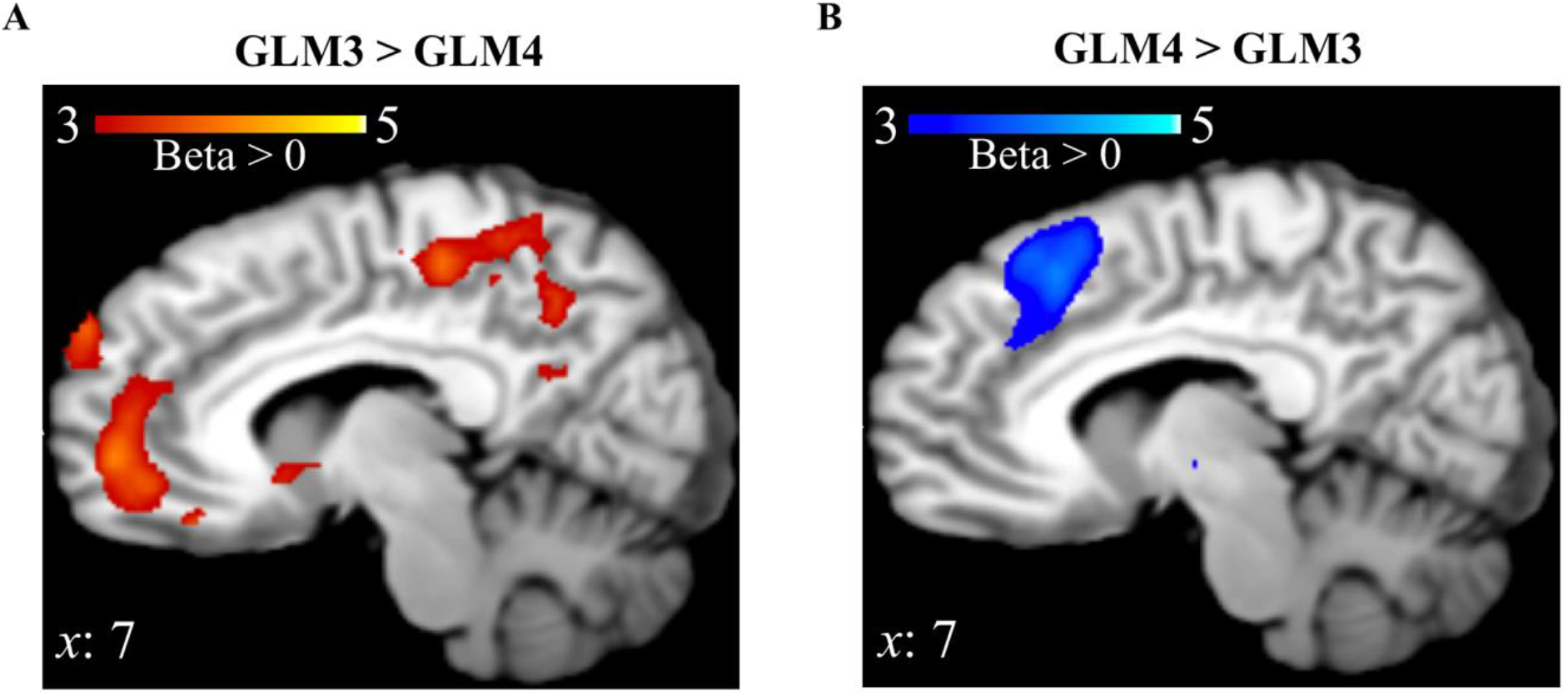
Domain specificity in vmPFC and dACC. A paired t-test between GLM3 and GLM4 shows **A**) specificity for reward (and not for information) in vmPFC, and **B**) for information (and not for reward in dACC).

**Table S1.** vmPFC and dACC opposition across different decision-making contexts.

| STUDIES | VMPFC | dACC |
| --- | --- | --- |
| Blair *et al.* 2006 | Greater reward | Lesser punishment |
| Kolling *et al.* 2012 | Value of engaging | Value of foraging |
| Boorman *et al.* 2013 | Immediate reward | Long-term reward |
| Chen *et al.* 2014 | Positive correlation with creativity questionnaire | Negative correlation with creativity questionnaire |
| Kolling *et al.* 2014 | Reduced risk | Increased risk |
| Skvortsova *et al.* 2014 | Expected reward | Negative Expected reward |
| Becker *et al.* 2016 | Exploit reward | Switch to alternatives |
| Sinha *et al.* 2016 | Early decrease, later increase during stress manipulation | Early increase, later decrease during stress manipulation |
| Shenhav *et al.* 2016 | Choice ease | Choice difficulty |
| Delaveau *et al.* 2017 | Default Mode Network | Task Positive Network |
| Arulpragasam *et al.* 2018 | Subjective value | Negative Subjective value |
| Suarez-Jimenez *et al.* 2018 | Learning danger | Approaching danger |

The table shows a selection of studies that report the symmetric opposition between vmPFC and dACC in value-based decision-making.

**Table S2.** Model estimated parameters from participants’ behavior.

|  | gkRL |  |  |  |
| --- | --- | --- | --- | --- |
| Participants | $\alpha$ | $\beta$ | $\omega$ | $\text{Log}(\gamma)$ |
| Number1 | 0.004 | 7.916 | 0.457 | -1.895 |
| Number2 | 0.151 | 0.443 | 2.723 | 0.098 |
| Number3 | 0.684 | 0.189 | 17.52 | -13.81 |
| Number4 | 0.495 | 0.358 | 33.333 | -1.61 |
| Number5 | 0.518 | 0.201 | 12.980 | -1.46 |
| Number6 | 0.459 | 0.168 | 48.663 | -13.81 |
| Number7 | 0.528 | 0.147 | 9.181 | -0.748 |
| Number8 | 0.593 | 0.158 | 24.743 | -1.84 |
| Number9 | 0.658 | 0.214 | 33.661 | -1.84 |
| Number10 | 0.054 | 1.819 | 1.714 | 0.138 |
| Number11 | 0.761 | 0.155 | 31.631 | -18.07 |
| Number12 | 0.687 | 0.142 | 1.114 | -0.003 |
| Number13 | 0.660 | 0.125 | 29.913 | -2.64 |
| Number14 | 0.393 | 0.321 | 17.714 | -19.94 |
| Number15 | 0.23 | 0.338 | 4.808 | -0.134 |
| Number16 | 0.269 | 0.791 | 7.528 | -0.425 |
| Number17 | 0.046 | 0.244 | 3.032 | -2.43 |
| Number18 | 0.269 | 0.208 | 9.174 | -0.178 |
| Number19 | 0.011 | 1.562 | 0.508 | -0.642 |
| Number20 | 0.012 | 2.084 | 0.26 | -0.080 |
| Total | 0.374<br>(0.264) | 0.879<br>(0.176) | 14.53<br>(1.76) | 4.07<br>(6.51) |

The table shows parameter estimates after fitting the model to participants' data. Group mean and standard deviation are also reported for each parameter.

**Table S3.** GLMs for fMRI data.

| *NAME* | *REGRESSORS* |
| --- | --- |
| GLM0 | [Highest Reward choice; Lower Reward choice] |
| GLM1 | [First free choice; $RQ_{t+1,j}(c)$; 24 motion regressors] |
| GLM1bis | [First free choice; $Q_{t+1,j}(c)$; 24 motion regressors] |
| GLM2 | [First free choice; $-I_{t,j}(c)$; 24 motion regressors] |
| GLM3 | [First free choice; $-I_{t,j}(c)$; $RQ_{t+1,j}(c)$; 24 motion regressors] |
| GLM4 | [ First free choice; $RQ_{t+1,j}(c)$; $I_{t,j}(c)$; 24 motion regressors] |
| GLM4bis | [ First free choice; $Q_{t+1,j}(c)$; $-I_{t,j}(c)$; 24 motion regressors] |
| GLM5 | [ First free choice; *Instrumental Info*($c$); $-I_{t,j}(c)$; 24 motion regressors] |
| GLM6 | [ First free choice; $-I_{t,j}(c)$; *Instrumental Info*($c$); 24 motion regressors] |
| GLM7 | [Defulat Option; Switch Option] |
| GLM8 | [ First free choice; $P(c/V_{t,j}(c_i))$; 24 motion regressors] |
| GLM9 | [ First free choice; $RQ_{t+1,j}(c)$; $Q_{t+1,j}(c)$; 24 motion regressors] |
| GLM10 | [ First free choice; $Q_{t+1,j}(c)$; $RQ_{t+1,j}(c)$; 24 motion regressors] |
| GLM11 | [ First free choice; $RQ_{t+1,j}(c)$; Max $Q_{t+1,j}$; Min $Q_{t+1,j}$; mean $Q_{t+1,j}$ ; 24 motion regressors] |

The table shows the 13 GLMs adopted in the fMRI data analysis all referring to activity associated with the onset of the first-free choice trial. GLM0 and 7 are the univariate analysis, whereas the other GLMs relate with the model-based analysis.

**Table S4.** Brain activity no reported in the text.

| GLM | ROI | Statistics<br>(cluster level<br>correction) |
| --- | --- | --- |
| GLM0<br>*Highest reward - Lower Reward* | PCC (8, -44, 28)<br>PCC (-6, -22, 42)<br>mOFC (-10, 60, 16) | $p < 0.001$<br>$p < 0.001$<br>$p = 0.037$ |
| GLM1 (Beta > 0) | PCC (-8, -50, 28) | $p < 0.001$ |
| GLM2 (Beta > 0) | right aInsula (34, 22, -8)<br>left aInsula (-34, 20, 6)<br>right dLPFC (42, 16, 40) | $p < 0.01$<br>$p < 0.05$<br>$p = 0.001$ |
| GLM2 (Beta < 0) | PCC (-10, -34, 46) | $p < 0.001$ |
| GLM3 (Beta > 0) | PCC (-2, -50, 26) | $p < 0.001$ |
| GLM4 (Beta > 0) | right aInsula (34, 22, -8 )<br>left aInsula (-38, 18, -10) | $p < 0.05$<br>$p < 0.05$ |
| GLM8 (Beta > 0) | PCC (-10, -32, 58) | $p < 0.002$ |

The table shows brain activity not reported in the main text. PCC: Posterior Cingulate Cortex; mOFC: medial Orbitofrontal Cortex; aInsula: anterior Insula

**Table S5.** Correlation of covariates with relative reward value.

| Subject | Max Value | Min Value | Standard Deviation | Averaged Value | Chosen-Second |
| --- | --- | --- | --- | --- | --- |
| 1 | 0.135 | 0.135 | 0.718 | -0.123 | 0.938 |
| 2 | 0.333 | 0.333 | 0.657 | -0.013 | 0.889 |
| 3 | 0.115 | 0.115 | 0.672 | -0.116 | 0.917 |
| 4 | 0.504 | 0.504 | 0.574 | -0.061 | 0.884 |
| 5 | 0.162 | 0.162 | 0.649 | -0.282 | 0.924 |
| 6 | 0.441 | 0.441 | 0.586 | -0.048 | 0.892 |
| 7 | -0.082 | -0.082 | 0.65 | -0.461 | 0.938 |
| 8 | 0.539 | 0.539 | 0.54 | 0.054 | 0.881 |
| 9 | 0.355 | 0.355 | 0.488 | -0.049 | 0.882 |
| 10 | 0.34 | 0.34 | 0.649 | -0.163 | 0.927 |
| 11 | -0.025 | -0.025 | 0.603 | -0.319 | 0.920 |
| 12 | 0.386 | 0.386 | 0.551 | -0.105 | 0.920 |
| 13 | 0.646 | 0.646 | 0.547 | 0.074 | 0.881 |
| 14 | 0.327 | 0.327 | 0.584 | -0.13 | 0.914 |
| 15 | 0.511 | 0.511 | 0.646 | 0.03 | 0.890 |
| 16 | 0.396 | 0.396 | 0.638 | -0.016 | 0.896 |
| 17 | 0.474 | 0.474 | 0.709 | 0.018 | 0.897 |
| 18 | 0.327 | 0.327 | 0.714 | 0.101 | 0.918 |
| 19 | 0.101 | 0.101 | 0.681 | -0.247 | 0.923 |
| 20 | 0.117 | 0.117 | 0.68 | -0.14 | 0.941 |
| 21 | 0.143 | 0.143 | 0.595 | -0.062 | 0.900 |

